# Reducing shade avoidance can improve Arabidopsis canopy performance against competitors

**DOI:** 10.1101/792283

**Authors:** Chrysoula K. Pantazopoulou, Franca J. Bongers, Ronald Pierik

**Affiliations:** Plant Ecophysiology, Dept. of Biology, Utrecht University, The Netherlands; State Key Laboratory of Vegetation and Environmental Change, Institute of Botany, Chinese Academy of Sciences, Beijing, China

**Keywords:** Shade avoidance, canopy architecture, planting pattern, competition, *Arabidopsis thaliana*, hyponasty

## Abstract

The loss of crop yield due to weeds is an urgent agricultural problem. Although herbicides are an effective way to control weeds, more sustainable solutions for weed management are desirable. It has been proposed that crop plants can communally suppress weeds by shading them out. Shade avoidance responses, such as upward leaf movement (hyponasty) and stem or petiole elongation, enhance light capture of individual plants, increasing their individual fitness. The shading capacity of the entire crop community might, however, be more effective if aspects of shade avoidance are suppressed. Testing this hypothesis in crops is hampered by the lack of well-characterized mutants. We therefore investigated if *Arabidopsis* competitive performance at the community level against invading competitors is affected by the ability to display shade avoidance. We tested two mutants: *pif4pif5* that has mildly reduced petiole elongation and hyponasty and *pif7* with normal elongation but absent hyponasty in response to shade. Although *pif4pif5* performed similar to wildtype, we found that *pif7* showed significantly increased canopy biomass and suppression of invading competitors as compared to its wildtype. Our data thus show that modifying specific shade avoidance aspects has potential for plant community performance. This may help to suppress weeds in crop stands.

**Highlight:** Hyponastic response in canopies facilitates light penetration and weed growth. Inhibition of this response to neighbors increased canopy biomass, canopy closure and suppression of competitors.

## Introduction

Competition from weeds accounts for substantial yield losses in global crop production systems (Bridges, 1994; Liebman *et al*., 2001; Oerke 2006). Weed control is usually accomplished by the extensive use of herbicides, and although these can be effective, they are costly and have negative side effects on people’s health and the environment (Buhler, 2002; Chauhan and Johnson, 2010). There is, therefore, an urgent need for novel methods to suppress weeds in a cost-effective and sustainable manner.

Light is the core energy source for all plants, and when grown at high planting densities, individual plants consolidate light capture by growing away from the shade cast by neighboring plants, a process called shade avoidance (de Wit *et al*., 2016*a*). Proximate plants are detected through the red (R) : far-red (FR) light ratio (R:FR) in the light reflected between plants, a ratio that decreases because of selective absorption of R light for photosynthesis and reflection of FR light. When weeds emerge in a crop vegetation, FR reflection becomes more intense due to the presence of more plant biomass, leading to strong reduction of R:FR (Pierik and Testerink, 2014). Several studies have shown that under low R:FR conditions, crops develop phenotypes such as inhibition of lateral branching (tillering) in wheat (Ugarte *et al*., 2010), formation of smaller tubers and longer stem in potato (Boccalandro *et al*., 2003) and increased stem elongation in maize (Dubois *et al*., 2010). All these morphological changes have a negative impact on crop productivity (Robson *et al*., 1996; Boccalandro *et al*., 2003).

One approach to improve crop yield and suppress weeds, is by optimizing the shading capacity of the entire crop community, for example by optimizing planting patterns and plant phenotypic responses to density. In Evolutionary Agroecology (a.k.a. Darwinian Agriculture) it is proposed that shade avoidance responses that enhance individual plant performance, actually reduce performance of the entire community of crop plants (Weiner *et al*., 2010).

Studies on cereal crops have shown that optimization of cropping pattern and density based on the community, rather than the individual performance, led to a more effective weed suppression and increased crop productivity (Weiner *et al*., 2001; Olsen *et al*., 2005, 2006; Kristensen *et al*., 2008). This has for example been shown to increase wheat yield by up to 30% (Weiner *et al*., 2001). This major improvement may be explained by the fact that crop plants growing in uniform patterns can collectively create a stronger shade over the weeds than those sown in for example row planting patterns. In other words, crop plants can collectively suppress weeds much better than individual plants would be capable of. Since shade avoidance responses have a major impact on the 3D architecture of the plant, they strongly impact the shading capacity of a uniform plant community. Here, we will investigate to what extent modulation of shade avoidance responses at high planting density can improve shading capacity, competitor (weed) suppression and canopy plant performance.

Shade avoidance responses are typically elicited upon detection of a reduced R:FR ratio, and are further promoted by depletion of blue light when the canopy closes ((Ballaré, 1999; de Wit *et al*., 2016*b*). Shade avoidance responses include upward leaf movement (hyponasty), elongation of stems and petioles and inhibition of branching (Franklin, 2008; Pierik and de Wit, 2013; de Wit *et al*., 2016*b*). These responses help plants reposition their leaves away from the shade and into the light. Shade avoidance responses have been observed in most crop species and also in a variety of wild species, including the genetic model plant *Arabidopsis thaliana* (Ballaré, 1999; Franklin, 2008; Martínez-García *et al*., 2010; Casal, 2012; Gommers *et al*., 2013).

In responses to low R:FR, phytochrome photoreceptors are inactivated (Ballaré, 1999; Franklin *et al*., 2003; Kozuka *et al*., 2010) and this relieves their repression of Phytochrome Interacting Factors (PIFs) (Li *et al*., 2012; Jeong and Choi, 2013; Leivar and Monte, 2014), a class of transcription factors that promote the expression of growth promoting genes (Oh *et al*., 2012; Zhang *et al*., 2013). PIF4, PIF5 and PIF7 are the dominant PIF proteins involved in shade avoidance in Arabidopsis (Lorrain *et al*., 2008; Koini *et al*., 2009; Hornitschek *et al*., 2012; Li *et al*., 2012; Pantazopoulou *et al*., 2017).

Shade avoidance responses in crops are often associated with reduced yield, because resource investments are rerouted from harvestable organs towards stem elongation (Robson *et al*., 1996; Boccalandro *et al*., 2003; Carriedo *et al*., 2016). These responses furthermore create a more open canopy architecture, which allows more light penetration that in turn facilitates weed growth.

Here, we will investigate if canopy planting patterns and modifications of shade avoidance responses can indeed optimize canopy architecture to suppress competitors by improved shading capacity. We will compare *Arabidopsis thaliana* wildtype with well-described mutants for aspects of the shade avoidance syndrome; an opportunity that does not (yet) exist in other plant species. We show that the *pif7* mutant that lacks a hyponastic response, but has preserved petiole elongation responses to neighbors, performs significantly better at high density, uniform planting patterns, than does its corresponding wildtype. Modest inhibition of both elongation and hyponasty in the *pif4pif5* double mutant on the other hand did not affect plant performance. Consistent with our hypothesis, a canopy of *pif7* plants was better able to suppress competing invaders than was a shade avoiding wildtype canopy.

Our data indicate that modifying plant canopy architecture through altered shade avoidance characteristics provides great opportunity to control weed proliferation in cropping systems in a sustainable way.

## Materials and methods

### Canopy conditions and measurements

Genotypes used in this study, as canopy plants were wild-type Col-0, *pif4-101 pif5-1* (Lorrain *et al*., 2008) and *pif7* (Leivar *et al*., 2008) while *pif4-101 pif5-1 pif7-1* (de Wit *et al*., 2015) was used as the invading competitor. Canopy seeds were sown in a pot with a surface area of 10.5×10.5 cm filled with a substrate of soil:perlite (2:1), with additional nutrients [6 g of slow release fertilizer (Osmocote ‘plus mini’ Ammonium Nitrate Based Fertilizer; UN2071; Scotts Europe BV, Heerlen, The Netherlands) and 6 g MgOCaO (17%; Vitasol BV, Stolwijk, The Netherlands]. The *pif4pif5pif7* plants were sown in a different pot three days after canopy plants for germination. Sowing was followed by stratification for four days (dark, 4°C). After stratification plants were moved to a short-day growth chamber (9 h/15 h of light/dark period respectively; R:FR was 2.3 and PAR = 150 µmol m^-2^ s^-1^). When canopy plots were 15 days old (seeds of the canopy were sown directly in plots), competitor *pif4pif5pif7* seedlings (12 days old) were transplanted into the plot (Fig. S1). The canopy plots were grown for another 29 days and subsequently harvested. Measurements were performed on four plants for each plot. Petiole and lamina length of the three longest leaves from each plant were measured with a digital caliper. Individual plant leaf area was scanned and determined with image-J software. Shoot dry weight was recorded with a digital scale, after drying the tissue at 70°C oven for three days. Plot biomass and LAI were calculated from the four individuals by extrapolating to the full plot and density. The heights of the canopies were measured with a ruler while the canopy cover was determined from top photographs using the Plant CV software (Gehan *et al*., 2017). In the canopy cover measurements by the PlantCV, we exclude the outer plants of the canopy to avoid edge effects. The height and the canopy cover measurements were taken every five days, starting from the day 20 of canopy growth. Seed output was recorded in separate experiments with the same growth conditions, three months after sowing. Every 10 days (starting from the sowing day) plants were watered with nutrients, on all other days they were watered with tap water. When the first silique from each pot turned brown, watering was stopped. The number of siliques was measured, after two weeks of ripening. Petiole angles (hyponasty) (Fig.3 and Fig.4) and lamina length (Fig. S5) of the fifth-youngest leaf were measured digitally with image J. Pictures were taken every day for 13 days, starting at day 28 (t=0).

### R:FR measurements

The R:FR measurements started at day 20 (before the competition starts, (de Wit *et al*., 2012)] by using the Spectrosense2-Skye light sensor with a glass fiber extension with 0.6 cm light collection area. The sensor was placed inside of the canopy plot (Fig. S1A) and measured the R:FR from four different directions and on four different positions, resulting in 16 measurements per time/per pot. When canopy closure occurred, the sensor was placed under the canopy, without causing any damage to the plants or interfering with the canopy shade. The measurements were always taken from the same position in all densities and patterns.

### Experimental design of the densities and patterns with or with the competitor *pif4pif5pif7*

For the Col-0 canopy plants three different densities were used (16 plants per pot (1111 plants m^-2^), 25 plants per pot (2500 plants m^-2^), 64 plants per pot (8264 plants m^-2^); hereafter low, medium and high density respectively) and two spatial patterns [uniform (equal distance between the plants) and row (bigger distance between the rows of the plants but smaller distance between the plants within the rows), See Fig. S1B)]. In uniform pattern, the distance between the plants was 3 cm, 2 cm and 1 cm in low, medium and high density respectively. In row pattern, the distance between the rows was always 5 cm while within the rows the distance between the plants were 0.6 cm, 1.25 cm and 2 cm in high medium and low density respectively. For the canopies consisting of *pif4pif5 and pif7* plants only high density-uniform pattern was used. The number of competitor *pif4pif5pif7* plants and their positions which were transplanted into the high density-uniform pattern plots were the same in all the canopies [16 plants per pot (1111 plants m^-2^)] (Fig. S1C).

### Light experiments

Individual plant responses to R:FR were studied for the different genotypes used here. To reduce the R:FR light ratios in the control white (W) light conditions from Philips HPI lamps (R:FR = 2.3, 160 μmol m^-2^ s^-1^ PAR), supplemental far-red LEDs (Philips Green Power FR 730 nm) were used. FR supplementation resulted in R:FR = 0.2 (160 μmol m^-2^ s^-1^ PAR). To mimic the true canopy shade, green filter (Lee filters Fern Green) was used (resulting in R:FR = 0.35 and 35 μmol m^-2^ s^-1^ PAR). The light spectra of the treatments were measured with an Ocean optics JAZ spectroradiometer (Fig. S2).

### FSP model

A functional-structural plant (FSP) model (Vos *et al*., 2009) of Arabidopsis rosettes, previously used and described in (Bongers *et al*., 2017, 2019; Pantazopoulou *et al*., 2017), was used to simulate Arabidopsis plant types, using the simulation platform GroIMP and its radiation model (https://sourceforge.net/projects/groimp/). Arabidopsis rosettes were represented by a collection of leaves (represented by a petiole and lamina) whose appearance rate and shape were based on empirical data (Bongers *et al*., 2017). The leaves individually grew in time in 3D based on light interception, photosynthesis and plant-wide carbon allocation principles (for detailed explanations of the principles see Evers, 2016; Bongers *et al*., 2017). In addition, leaves showed petiole elongation and hyponastic responses based on the virtual touching of leaves and the perception of R:FR (Bongers *et al*., 2017). Therefore, individual growth and shade avoidance responses depended on the capture of light (represented by PAR intensity) and the perception of R:FR within the simulated canopy.

The light source emitted PAR representing 220 μmol m^-2^ s^-1^ and a R:FR ratio of 2.3, which corresponded to the growth chamber experiments. 100 plants were placed in a uniform grid of 10 × 10 with an inter-plant distance of 1 cm, of which only the middle 16 plants were used for analyses. Plants grew for 44 days based on the PAR captured, photosynthesis rates and carbon allocation patterns (Evers, 2016; Bongers *et al*., 2017). Each model time step, which represented 24 hours, hyponastic responses could occur if leaves touched or if R:FR perception at the lamina tip was below 0.5 (Bongers *et al*., 2017; Pantazopoulou *et al*., 2017). The strength of the hyponastic responses depended per model scenario; plants could increase their leaf angle with 0, 0.2, 1, 5, 10, 15 or 20 degrees per day. The angle of the leaves over time was therefore a function of the number of times in which touch and/or low R:FR perception occurred per individual leaf, with a maximum leaf angle of 80 degrees. The intensity of PAR that reached the soil was captured by 64 virtual soil-tiles (each 0.25 cm^2^) underneath the 16 middle plants.

### Statistics

Data were analyzed by one or two-way ANOVA followed by LSD test. All the analyses were performed with GraphPad.

## Results

### The effect of planting density and pattern on Col-0 performance

To investigate the effect of sowing pattern and density on *Arabidopsis thaliana* (hereafter Col-0) performance, we grew canopy plots in three different densities (low, medium and high) and two different patterns (uniform and row) (Fig. 1A). The R:FR showed a reduction in all densities and patterns through time, reflecting the growing canopy (Fig. 1B). However, the strongest and most rapid decline of R:FR was observed in high density/uniform pattern, where the R:FR was decreased from approximately 2.0 to 1.1 after eight days of measurements hinting at a rapidly closing canopy (Fig. 1A). This was not the case for the row pattern in high density, where the R:FR was still high, presumably because the inter row distance was higher than in the uniform pattern. Low and medium density showed reduction of R:FR (less than 1.5) at day 36 (Fig. 1B), indicating that the canopy remained more open for a longer period of time. The leaf area index (LAI) expresses the amount of leaf area per unit soil area and reflects the closure status of the canopy. LAI increased more strongly in the uniform than in the row pattern and mostly in the medium and high densities (Fig. 1C). Interestingly, leaf lamina length decreased with increasing plant density, irrespective of the planting pattern (Fig. S3A). The opposite was observed for petiole length, where the high density induced the strongest elongation (Fig. S3B). Enhanced petiole elongation, combined with reduced lamina size, are classic aspects of shade avoidance.

**Fig. 1:**
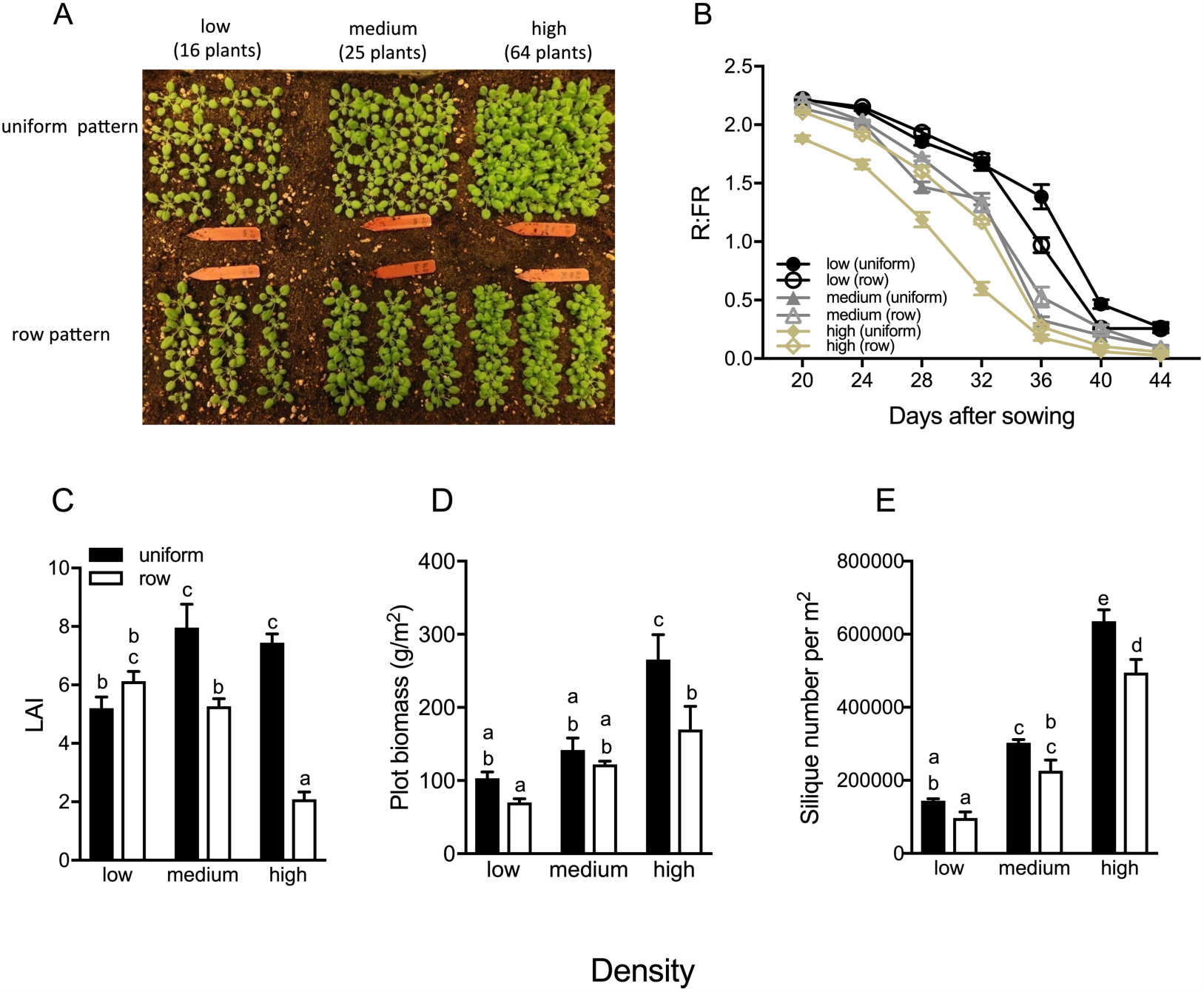
Arabidopsis Col-0 in high density, uniform pattern produces more biomass and canopy cover than at lower densities and row patterns. (A) In the upper row Col-0 plants grow in a uniform pattern (uniform), while in the lower row plants grow in row pattern (row) at three different densities (low, medium, high). (B) The R:FR light ratio measured inside Col-0 canopies, during the days of growth, in low (black lines), medium (grey lines) and high (yellow lines) densities and two patterns (uniform and row). (C-E) Leaf area index (LAI; C), plot biomass per m^2^ (D) and seed output (silique number per m^2^ pot; E) at three different densities (low, medium, high) and two different planting patterns (uniform, row). Data represent mean ± SE (n=5). Different letters indicate statistically significant differences (two-way ANOVA with LSD test, *P* < 0.05).

Furthermore, there was a strong and significant effect of the density and planting pattern on Col-0 biomass. The row pattern produced Col-0 plants with the smallest dry weight, indicating that the intraspecific competition was higher in rows compared to the uniform pattern. In terms of planting density, the total biomass of the plot in high density and uniform pattern was higher than the other densities (medium, low) than the row pattern (Fig. 1D).

The number of siliques per square meter for the different density and planting patterns was consistent with biomass (Fig. 1E), which suggests that the uniform-planting pattern at the high density would result in the highest yield per unit area.

### Steering canopy light penetration through variations in hyponasty

Using a previously published 3D computational Arabidopsis model (Bongers *et al*., 2017), we determined the percentage of light penetration in the canopy through time and under control of different degrees of hyponasty. Seven different hyponastic scenarios were simulated; from 0 degrees up to 20 degrees hyponastic growth (Fig. 2). The simulations show that canopies, consisting of plants with minimal hyponastic response to neighbors (e.g. 0,0.2, 1 and 5 degrees) create strong reduction of light penetration inside the canopy; after 28 days, less than 5 % of the light reaches the soil. On the other hand, scenarios with faster hyponasty allowed for less light extinction and thus higher penetration of light inside the canopy. These simulations support the notion that upward leaf movement responses to neighbors may facilitate light penetration through the canopy, which can be beneficial for weed growth.

**Fig. 2:**
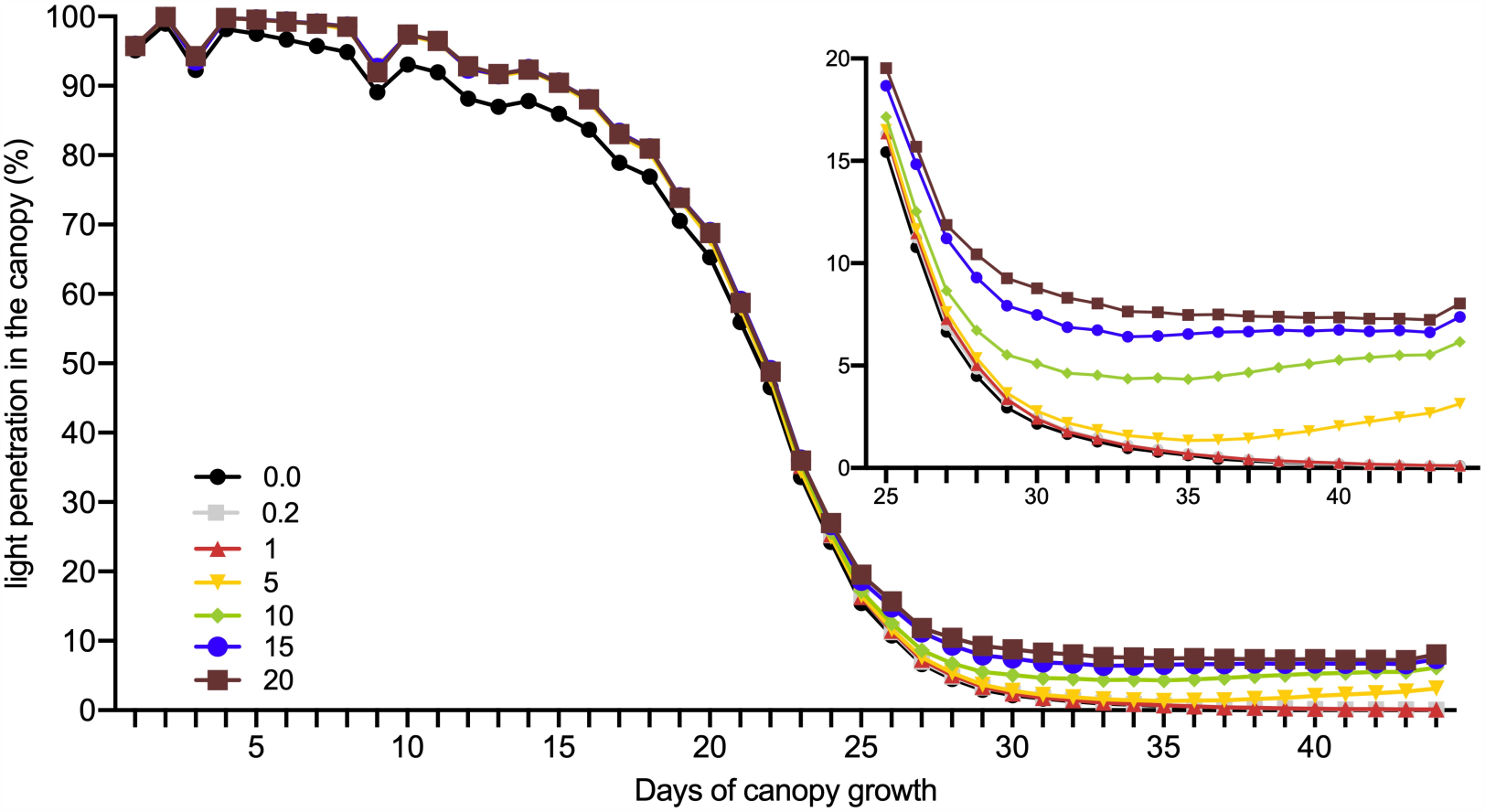
Reduced hyponastic responses result in lower percentage of light penetrating the canopy and reaching the soil. Percentage of light reaching the soil is simulated with a 3D Arabidopsis computational model. Various canopy growth simulation scenarios consist of plants with different degrees of hyponastic responses to proximate neighbor plants, ranging from 0 to 20 degrees per day (see legend). Data represent mean ± SD (n=10).

To test experimentally if altered hyponastic growth can regulate canopy closure and light penetration, we selected previously published mutants with altered shade avoidance characteristics. The *pif7, pif4pif5* double and *pif4pif5pif7* triple knockouts have reduced hyponastic responses to shade cues in short-term experiments (Pantazopoulou *et al*., 2017) and we verified their responses to prolonged shade cue conditions. Reduction in R:FR resulted in the elevation of Col-0 petiole angle (hyponasty) during the first two days (day 29 and 30), while petiole elongation was promoted from day 28 until 34 (Fig. 3A-D). *pif7* had a similar petiole elongation response as did Col-0 in all the treatments but its hyponastic response to low R:FR was entirely absent, whereas its response to green shade (reproducing canopy shade) was severely reduced (Fig. 3A & 3B). *pif4pif5* showed a phenotype initially similar to wild-type both in terms of petiole angle and elongation, but the petiole elongated slightly less through time in low R:FR. On the other hand, *pif4pif5pif7* was unresponsive to low R:FR for both traits (Figure 3C & 3D). Green filter triggered a continuous shade avoidance phenotype in Col-0 (hyponasty and petiole elongation) from day 28 up to 36 (8 days) (Figure 3B & 3D). Hyponastic responses were reduced in *pif4pif5* and not observed at all in *pif7 and pif4pif5pif7* under these severe shade conditions (Fig. 3A & 3C). In general, Col-0 shade avoidance responses (hyponasty & petiole elongation) were stronger in green shade than in low R:FR alone. Overall, *pif4pif5* was less responsive than Col-0, whereas *pif4pif5pif7* was fully insensitive to the different light conditions. Interestingly, *pif7* showed similar petiole growth as Col-0 and a similarly absent hyponastic response as in *pif4pif5pif7*.

**Fig. 3:**
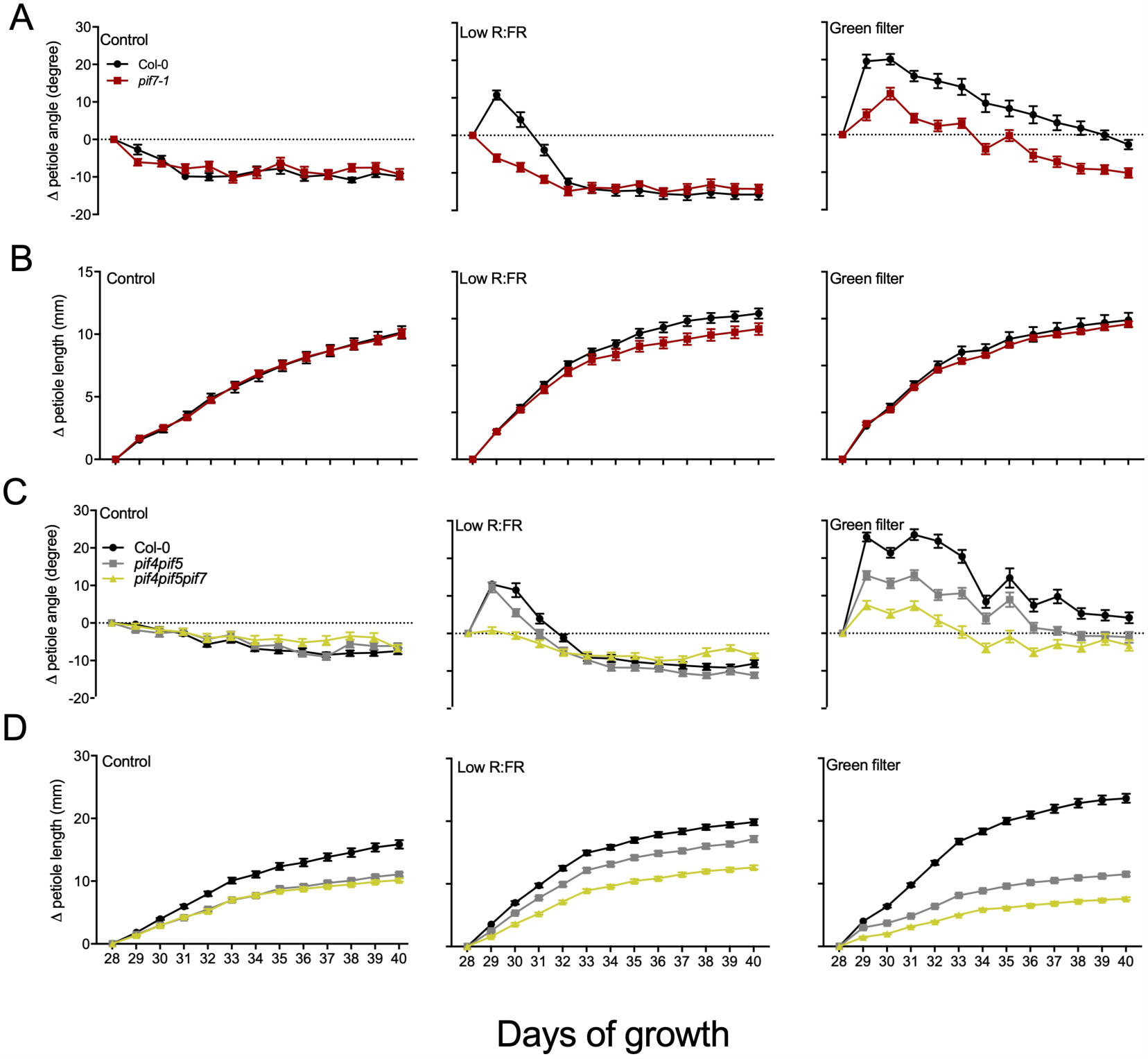
Shade avoidance responses (change in petiole length (A, C) and change in petiole angle (B, D)) of Col-0, *pif7, pif4pif5* and *pif4pif5pif7* upon white light (control), low R:FR and green filter exposure. Light treatments lasted 13 days and started when plants were 28 days old. Data represent mean ± SE (n=15).

The impact of different magnitudes of hyponastic responses in canopy closure, was tested by growing canopies of Col-0, *pif7* and *pif4pif5*. We decided not to use the *pif4pif5pif7* triple mutant as a canopy plant, but as an invading competitor in establishing canopies. High density, uniform planting patterns were used, since these closed their canopies most effectively (Fig. 1). Here, we monitored the canopy closure state through time by using the imaging analysis PlantCV (Fig. 4). Data showed that *pif7* canopies developed a better soil cover than Col-0 and *pif4pif5* early in the canopy development (day 20 until 25). The *pif4pif5* canopies remained more open than *pif7* canopies for another five days but percentage of the covered soil area was not significantly different from Col-0 canopies at day 30. At later stages all canopies had developed nearly full closure without. The *pif4pif5* canopies display reduced petiole elongation compared to *pif7* and Col-0 (Fig. 3D), resulting in a relatively low canopy height for this double mutant (Fig. S4B). The height of *pif7* canopies was also reduced as compared to Col-0 (Fig. S4B), presumably because of the reduced upward leaf movement in this mutant (Fig. 3A)

**Fig. 4:**
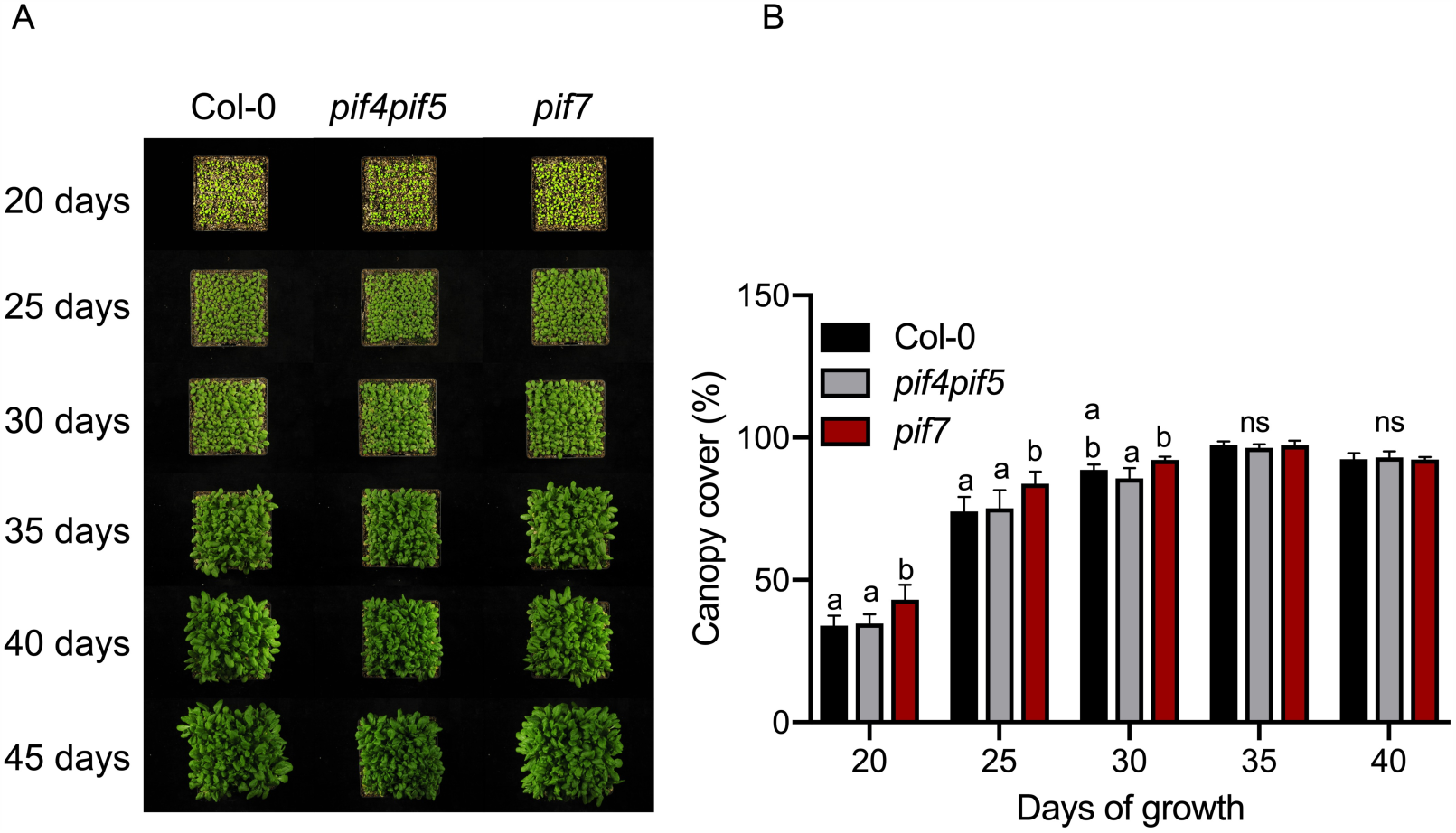
The *pif7* mutant creates a faster closed canopy than Col-0 and *pif4pif5*. (A) Pictures illustrate how the canopies of Col-0 (left), *pif4pif5* (middle) and *pif7* (right) plants develop and close soil exposure to light. (B) The percentage of soil covered by the same canopies: Col-0 (black bars), *pif4pif5* (grey bars) and *pif7* (red bars) plants, through time. The Col-0, *pif4pif5* and *pif7* canopies plants grew at high density, uniform pattern. Data represent mean ± SE (n=5). Different letters indicate statistically significant differences (two-way ANOVA with LSD test, *P* < 0.05. ns=not significant).

### Performance of canopy and competitor plants during competition

To test the impact of separate shade avoidance traits on competitor suppression and canopy performance, we used as canopy plants the strong shade avoider Col-0, the mild reduction of shade avoidance genotype *pif4pif5*, and *pif7* which does not show hyponasty but does induce petiole elongation upon low R:FR (Fig. 3). As an invading competitor we used *pif4pif5pif7*, which was planted between the canopy plants. To estimate shade avoidance responses of the different genotypes in true canopies, rather than independent light treatments, we measured petiole and lamina length at the end of the canopy development. *pif7* canopy plants displayed the largest lamina compared to Col-0 and *pif4pif5* canopy plants during competition. Petiole length was enhanced upon competition in Col-0 and *pif7* but not in *pif4pif5* canopy plants (Fig. S6). The strong lamina and petiole elongation but not hyponasty (fig. 3 and fig S4B) responses of *pif7* during competition could have resulted in the higher biomass and LAI compared to the other two genotypes (Fig. 5). This also had a strong effect on *pif4pif5pif7* competitor performance. The faster closed canopy and plant growth of *pif7* during competition was associated with a reduction in growth of *pif4pif5pif7* competitors (Fig. 6A & 6B). On the other hand, the improved light exposure of *pif4pif5pif7* competitor plants under the rapidly closed canopy of *pif4pif5* was associated with enhanced biomass and leaf area (L.A.) of the competitor triple mutant compared to the other genotypes (Fig. 5, 6A & 6B). Indeed, the *pif4pif5pif7* competitor hardly survives under the *pif7* canopy while the percentage of survival between Col-0 and *pif4pif5* was similar (Fig. 6C).

**Fig. 5:**
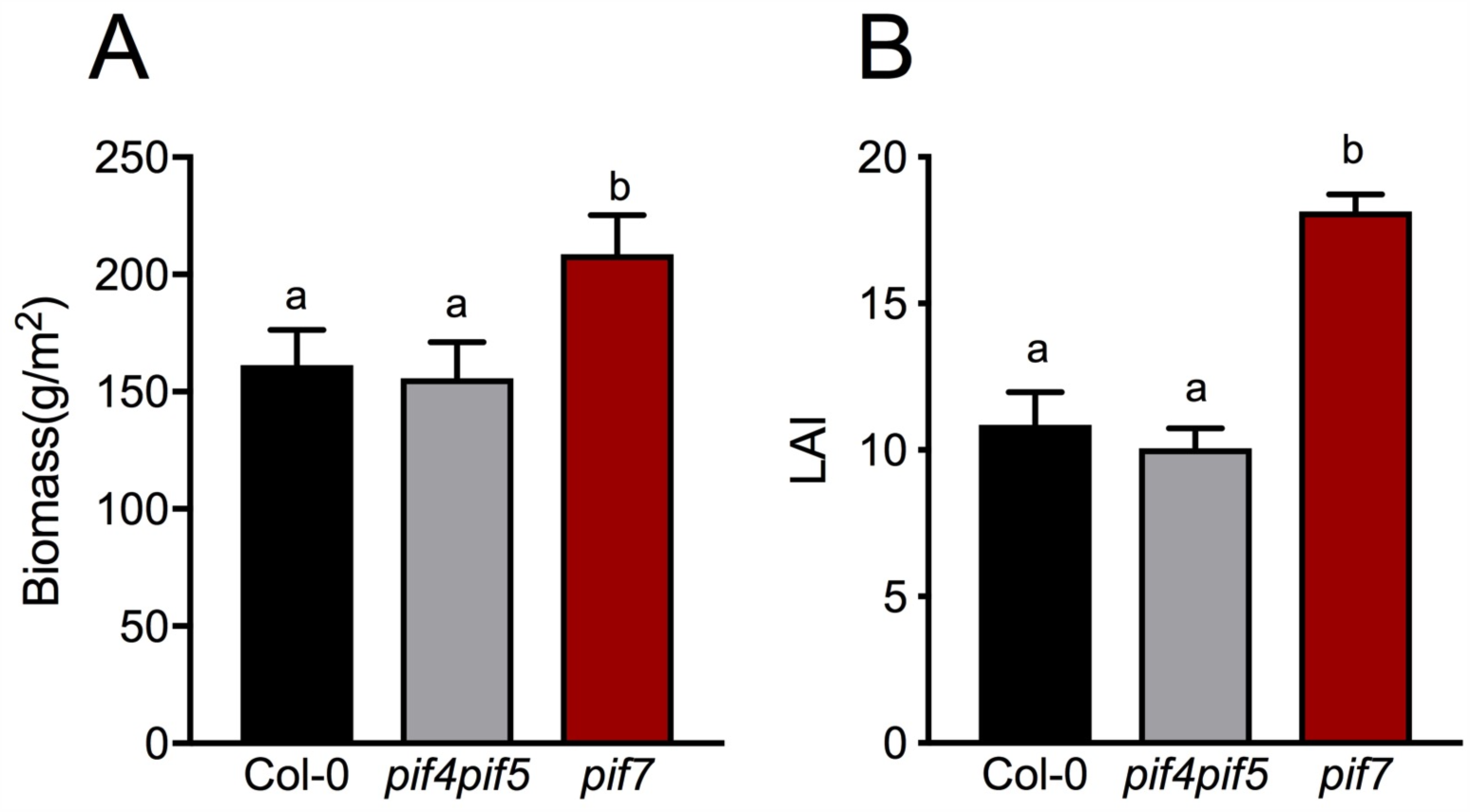
The *pif7* canopies grew larger than Col-0 and *pif4pif5* under high-density competing conditions. (A) Biomass and (B) LAI of canopies consisting of Col-0 (black bar), *pif4pif5* (grey bars) or *pif7* (red bars), growing at high density, uniform pattern, measured after 44 days of growth. Data represent mean ± SE (n=5). Different letters indicate statistically significant differences (two-way ANOVA with LSD test, *P* < 0.05).

**Fig. 6:**
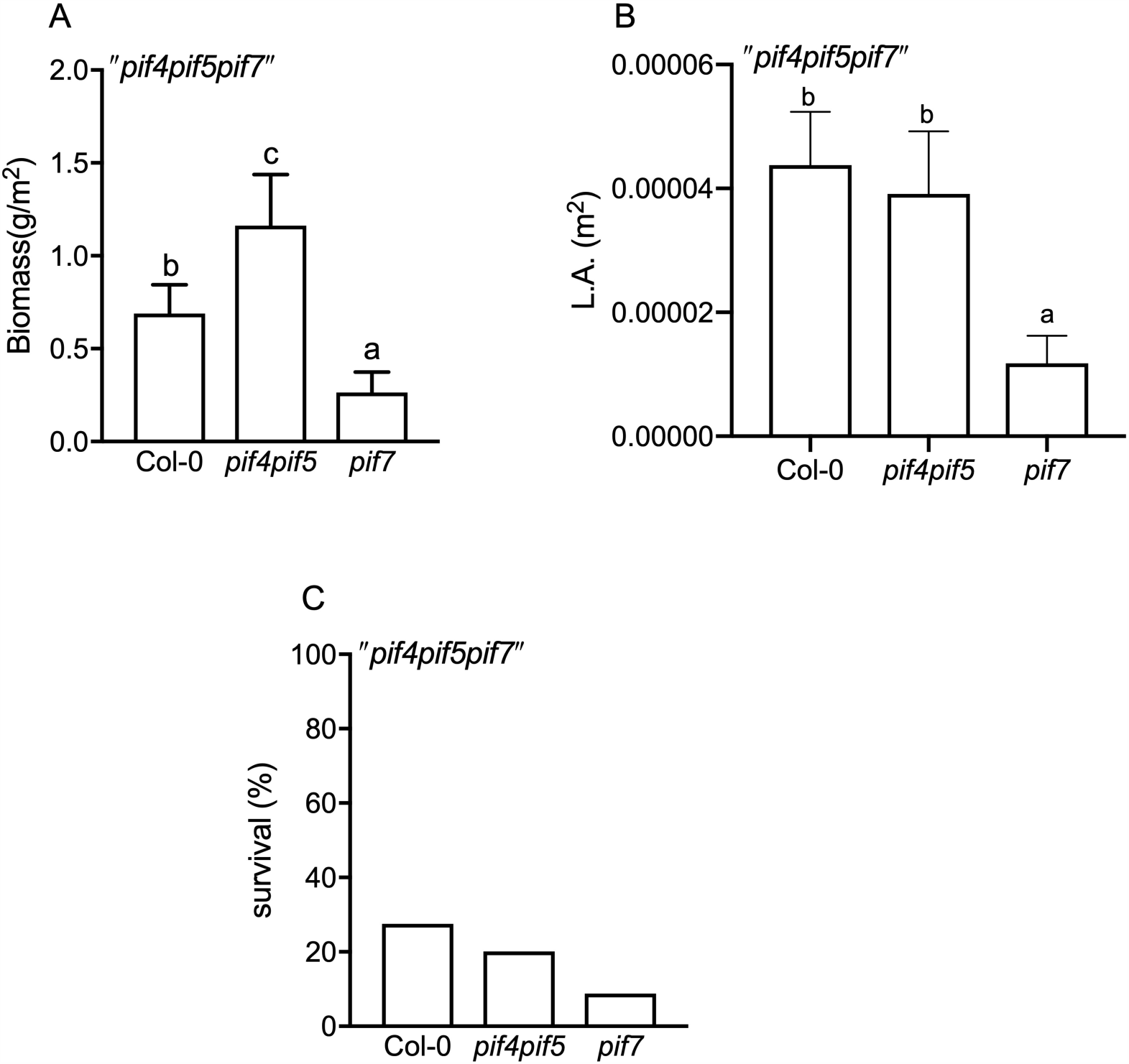
The *pif7* canopies suppressed the competitor*pif4pif5pif7* more effectively than did Col-0 and *pif4pif5* canopies. The competitor’s (A) biomass, (B) leaf area and (C) percentage of survival, under the canopies of Col-0, *pif4pif5* and *pif7* for 44days. The plants grew at high density, uniform pattern. Data represent mean ± SE (n=5). Different letters indicate statistically significant differences (two-way ANOVA with LSD test, *P* < 0.05).

## Discussion

The adaptability of to changing environments and their competitive potential makes them a major threat for agricultural yields when they interact with crop plants (Oerke, 2006). Studies in the past decades found that crop sowing uniformity in high density can positively affect yield and suppress weeds (Weiner *et al*., 2001; Olsen *et al*., 2006; Kristensen *et al*., 2008). A second way to improve crop yield and suppress weeds, according to the principles of Evolutionary Agroecology would be by controlling the shade avoidance properties of the crops in a way that would minimize light penetration through the canopy down to the soil where weeds sprout (Weiner *et al*., 2010). Since only very few well-defined shade avoidance mutants exist in crops (Carriedo *et al*., 2016; Kebrom and Mullet, 2016; Weiner *et al*., 2017), we tested the impact of shade avoidance modulation on canopy performance and weed suppression in the model species *Arabidopsis thaliana*.

Although petiole elongation, combined with upward leaf movement (hyponasty), will increase access to light at the individual plant level (Ballaré and Pierik, 2017; Pantazopoulou *et al*., 2017), the reduced leaf lamina growth that typically occurs in shade avoiding *Arabidopsis* (de Wit *et al*., 2015) may counterbalance the predicted gain in photosynthesis of individual plants (Fritz *et al*., 2018). Part of the shade avoidance responses will have been triggered through the drop in R:FR inside the canopies (Fig. 1B). However, shade avoidance responses, and especially hyponasty, can on their turn also affect the R:FR inside the canopy by affecting the extent to which a vertical canopy structure is formed in this otherwise horizontally growing rosette species (de Wit *et al*., 2012). Modulating shade avoidance traits in different canopy structures may thus affect light distribution inside these canopies. Indeed, using a 3D Arabidopsis plant model (Bongers *et al*., 2017), we found that slow-down of hyponastic growth upon shade detection in all canopy plants can clearly reduce light penetration through the canopy down to soil level (Fig. 2). To test the consequences of this scenario experimentally and also monitor the effect that these canopies can have on competitor performance we used *Arabidopsis* mutants that had similar hyponastic responses variations as used in the 3D plant model. As a competitor, we used the *pif4 pif5 pif7* triple mutant, which remained unresponsive in terms of hyponasty and petiole elongation under the long-term shade conditions. The *pif4pif5pif7* would not be able to outgrow the canopy plants, allowing us to 1) mimic the crop-weed competition where crops (like cereals) typically have a size advantage and 2) record the effect of different canopy architectures (i.e. *pif7* and *pif4pif5*) on competitor performance. The canopy architecture of *pif7* had a strong negative impact on the performance of competitor *pif4pif5pif7* and a positive impact on its own canopy biomass (fig. 5A). Under these conditions, the *pif4pif5pif7* competitor biomass and survival rate were significantly lower than under Col-0 and *pif4pif5* canopy architectures (Fig. 6A & 6C). We propose that the much faster closing of the *pif7* canopy together with the larger LAI as compared to the Col-0 and *pif4pif5* canopies (Fig. 4, S4 & fig. 5B), resulted in less light availability for the competitor, leading to reduced performance of the competitor.

Interestingly, despite the fact that the *pif4pif5* canopy architecture showed mild reduction of shade avoidance responses, the competitor *pif4pif5pif7* performed similar in Col-0 and *pif4pif5* canopy. We speculate that the advantage of modestly reduced shade avoidance in *pif4pif5* for communal competitor suppression might be outweighed by its reduced overall growth rate, which still leads to a relatively open canopy.

As mentioned above, *pif4pif5pif7* lacks any shade avoidance response to plant density, signaled by low R:FR and green filter (fig. 3). This allowed us to study if the resident canopy architecture can be optimized such that growth in the understory can be inhibited by shading the invading competitors. Future studies could be designed to include competitors that can show shade avoidance responses and thus have the capacity to compete stronger against the dominant canopy. It would be possible then that the invading competitors could even escape from the shade-casting canopy altogether and enhance their individual fitness at the expense of the collective fitness of the dominant canopy. If good mutants come available for upright-growing, stem-forming plants, these could be used to test scenario’s of a more vertically layered canopy, representing many of the staple crops world-wide, for weed-suppression.

Our data show that losing one of the shade avoidance responses in Arabidopsis canopy, hyponasty, has potential to suppress competitors. Translating this to crop-weed competition scenario’s depends on the architecture of the crop plant but could potentially improve weed suppression and reduce yield losses due to weeds.

## Author contributions

C.K.P. and R.P. designed research; C.K.P. and F.J.B. performed research; C.K.P. and F.J.B analyzed data; and C.K.P. and R.P. wrote the paper.

The authors declare no conflict of interest.

## Acknowledgments

We thank Tom Rankenberg for instructions on using PlantCV, Maxime Brugman for help with preliminary experiments and the Plant Ecophysiology group for help during harvests. This work was funded by Netherlands Organization for Scientific Research Vidi Grant 86512.003 (to R.P.)..

